# Peroxiredoxin 1 safeguards the nucleolar genome from oxidative damage

**DOI:** 10.1101/2025.11.12.685177

**Authors:** Takashi Furusawa, Vaibhavi Gujar, Shalu Sharma, Frimpong Boateng, Haojian Li, Cyrinne Achour, Daiki Taniyama, Yves Pommier, Shalini Oberdoerffer, Travis H. Stracker, Urbain Weyemi

**Author notes:** **Corresponding author:** Developmental Therapeutics Branch, National Cancer Institute, National Institutes of Health 37 Convent Drive, Bethesda, MD 20892, USA. These authors contributed equally to this study (Takashi Furusawa, Vaibhavi Gujar).

## Abstract

Peroxiredoxin 1 (PRDX1) is a highly conserved, thiol-dependent peroxidase that rapidly scavenges reactive oxygen species to modulate redox signaling. PRDX1-null mice exhibited genomic instability, shortened lifespan, and accelerated tumorigenesis, including development of lymphomas, sarcomas, and carcinomas. Despite extensive characterization of these phenotypes, the molecular mechanism by which PRDX1 loss causes genomic instability remains poorly understood. Here we show that PRDX1 deficiency alters nucleolar morphology, impairs RNA Polymerase I (POL-I)-dependent transcription of pre-ribosomal RNAs and triggers nucleolar genomic instability. This oxidative stress-induced nucleolar dysfunction promotes the stability of secondary DNA structures, such as RNA-DNA hybrids and G-quadruplex DNA, contributing to nucleolar genomic instability. We demonstrate that PRDX1 loss reduces nascent rRNA levels and impairs rRNA processing, further affecting ribosome biogenesis. Mechanistically, we established that PRDX1 loss triggers activation of the nucleolar DNA damage response including activation of DNA repair kinase ATM and the nucleolar factor TCOF1 within the nucleolus, and recruitment of the MRE11-RAD50-NBS1 (MRN) complex subunit NBS1 to ribosomal DNA (rDNA) loci. NBS1 accumulation correlates with the repression of rDNA transcription by POL-I, potentially delaying rRNA synthesis, and safeguarding the nucleolar genome from further oxidative damage. Collectively, these findings uncover a previously unrecognized, but critical role, for PRDX1 in maintaining nucleolar integrity and ribosomal biogenesis through redox-dependent regulation of rDNA transcription and processing machinery.

## Introduction

The nucleolus is a dynamic, membrane-less nuclear compartment that hosts the ribosomal DNA (rDNA) sequences essential for ribosome biogenesis ^1^. Roughly 30% of nucleolar proteins are dedicated to ribosome production, underscoring the central role of the nucleolar genome in coordinating diverse cellular functions^2^. The nucleolus is the primary site of RNA Polymerase I (POL-I)-mediated transcription of 45S/47S ribosomal RNA (rRNA) precursors, which are subsequently processed into 28S, 18S, and 5.8S rRNA. These mature rRNA molecules assemble with ribosomal proteins to form pre-ribosomes that mature into the 40S and 60S ribosomal subunits in the cytosol, where they drive protein synthesis ^1,3,4^. Because POL-I accounts for most of the cellular transcriptional activity, its dysregulation contributes to diverse pathologies, including cancers and ribosomopathies ^5^. Since the nucleolus may be exposed to genotoxic agents, it is endowed with robust and rapid sensors that prevent DNA lesions, including oxidant-induced DNA damage, safeguarding the nucleolar genome.

The MRN (MRE11-RAD50-NBS1) complex is a critical component of the DNA damage response (DDR) that recognizes DNA double strand breaks (DSBs), regulates DNA metabolism during repair and activates the ATM kinase, a central transducer of the DDR ^6^. However, it remains unclear whether ATM-dependent signaling cascades regulate rDNA transcription. Recent reports suggested that ATM-dependent phosphorylation of the nucleolar factor TCOF1 (Treacle) is sufficient to repress local POL-I-mediated transcription in the nucleolous following chromatin breaks ^7,8^. Another report demonstrated that POL-I activation is mediated by nucleolar recruitment of NBS1 and subsequent binding of the MRN complex to rDNA, implicating a role for NBS1 in ribosome biogenesis ^9^. Similarly, other findings identified TCOF1 as a DDR factor that cooperates with ATM and NBS1 to suppress inappropriate rDNA transcription and maintain genomic integrity after DNA damage ^10^.

Peroxiredoxins are thiol-dependent peroxidases that rapidly detoxify hydroperoxides and modulate redox signaling. Peroxiredoxins may transfer oxidative equivalents to partner proteins, or safeguard partners from oxidation-induced inactivation ^11–13^. PRDX1, a typical 2-Cys peroxiredoxin, reduces hydrogen peroxide (H_2_O_2_) or alkyl peroxide (ROOH) via its conserved peroxidatic cysteine residue Cys 52, which forms an intermolecular disulfide with the resolving cysteine Cys 173 of the partner subunit. This disulfide bond is subsequently reduced by the Thioredoxin (Trx)-Thioredoxin reductase (TrxR) complex and NADPH system, enabling continuous detoxification of peroxides ^14^. Mutation of both the peroxidatic and resolving cysteine residues disrupts disulfide formation and significantly abolishes peroxidatic activity. This redox cycle enables PRDX1 to continuously detoxify peroxides and relay oxidative equivalents. Upon elevated oxidation, PRDX1 can also undergo hyperoxidation or oligomerization, generating a chaperone complex that protects and stabilizes partner proteins ^14^.

Mounting evidence implicates PRDX1 in genomic stability, including seminal findings that demonstrated genomic instability, shortened lifespan, and accelerated tumorigenesis, including development of lymphoma, sarcomas, and carcinomas, in PRDX1-null mice ^15^. In addition, PRDX1 was shown to interact with DDR proteins to promote genomic stability, including RAD51 that was protected from stress induced oxidation by PRDX1 ^16–18^. Recent reports also linked PRDX1 to additional genome maintenance functions, including protecting telomeres from oxidation ^19^ and maintaining nucleotide pool synthesis following exposure to replication stress ^20^. We previously demonstrated that PRDX1 loss induced genomic instability and conferred synthetic lethality with DDR inhibitors, such as ATR and ATM inhibitors ^21^. Although PRDX1 localizes to both the cytosol and nucleus, the extent to which its nuclear localization contributes to genomic stability and RNA metabolism remains unclear.

Here, we uncover an essential nucleolar function for PRDX1 in genome maintenance. Using sub-cellular fractionation followed by measurement of markers of nucleolar genomic instability, we show that PRDX1 is required for nucleolar homeostasis. Loss of PRDX1 alters nucleolar morphology and impairs POL-I-mediated rDNA transcription, rRNA synthesis and subsequent processing steps, culminating in disruptive ribosome biogenesis. We demonstrated that PRDX1 deficiency is accompanied by DDR activation in the nucleolus including ATM activation and nucleolar accumulation of the nucleolar factor TCOF1, together with recruitment of the MRN subunit NBS1 to rDNA loci upon oxidative stress. This nucleolar DDR is associated with limited rDNA transcription, potentially preserving the nucleolar genome from oxidative stress-induced genomic instability. Collectively, these findings reveal PRDX1 as a key redox regulator of nucleolar homeostasis and rDNA integrity.

## Results

### PRDX1 loss is associated with increased frequency of cells with nucleolar abnormalies

While the cytosolic and nuclear localization of PRDX1 has been widely characterized ^11,22,23^, its nuclear function remains poorly understood, despite studies linking PRDX1 to genomic stability ^16,19,20,24^. To explore the nuclear role of PRDX1, we overexpressed it in the human non-small cell lung cancer line H1299, and carried out a cell fractionation assay. This revealed that while a large fraction of PRDX1 is largely distributed to the cytoplasm, it was also found to be enriched in the nucleoplasm and enriched nucleolar fraction to similar levels (Fig. 1A). As previous studies have suggested that loss of proteins that colocalize to the nucleolus have significant impact on the nucleolar morphology, these observations prompted us to determine the extent to which PRDX1 loss or H_2_O_2_ treatment affected nucleolar morphology by comparing parental and PRDX1 knockout cells stained with the nucleolar marker Nucleolin (NCL) and analyzed using confocal imaging (Fig. 1B). More than 75% of parental A549 lung cells exhibited nucleoli with irregular borders, as revealed by the distribution of the canonical nucleolar marker Nucleolin (NCL) (Fig. 1C and 1D) ^1^. In contrast, PRDX1 loss led to a 50% reduction in cells harboring the canonical nucleolar shapes, while the rest of the PRDX1 knockout cells showed more round and regular-shaped nucleoli (Fig. 1C and 1D), with a few cells exhibiting peripheral nucleolar distribution of NCL, known as “caps-like” nucleoli (Fig. 1E). H_2_O_2_ treatment partially mimics PRDX1 loss. These findings suggest that abnormalities in nucleolar morphology result from PRDX1 depletion and point to PRDX1 as key a determinant of nucleolar homeostasis.

**Figure 1:**
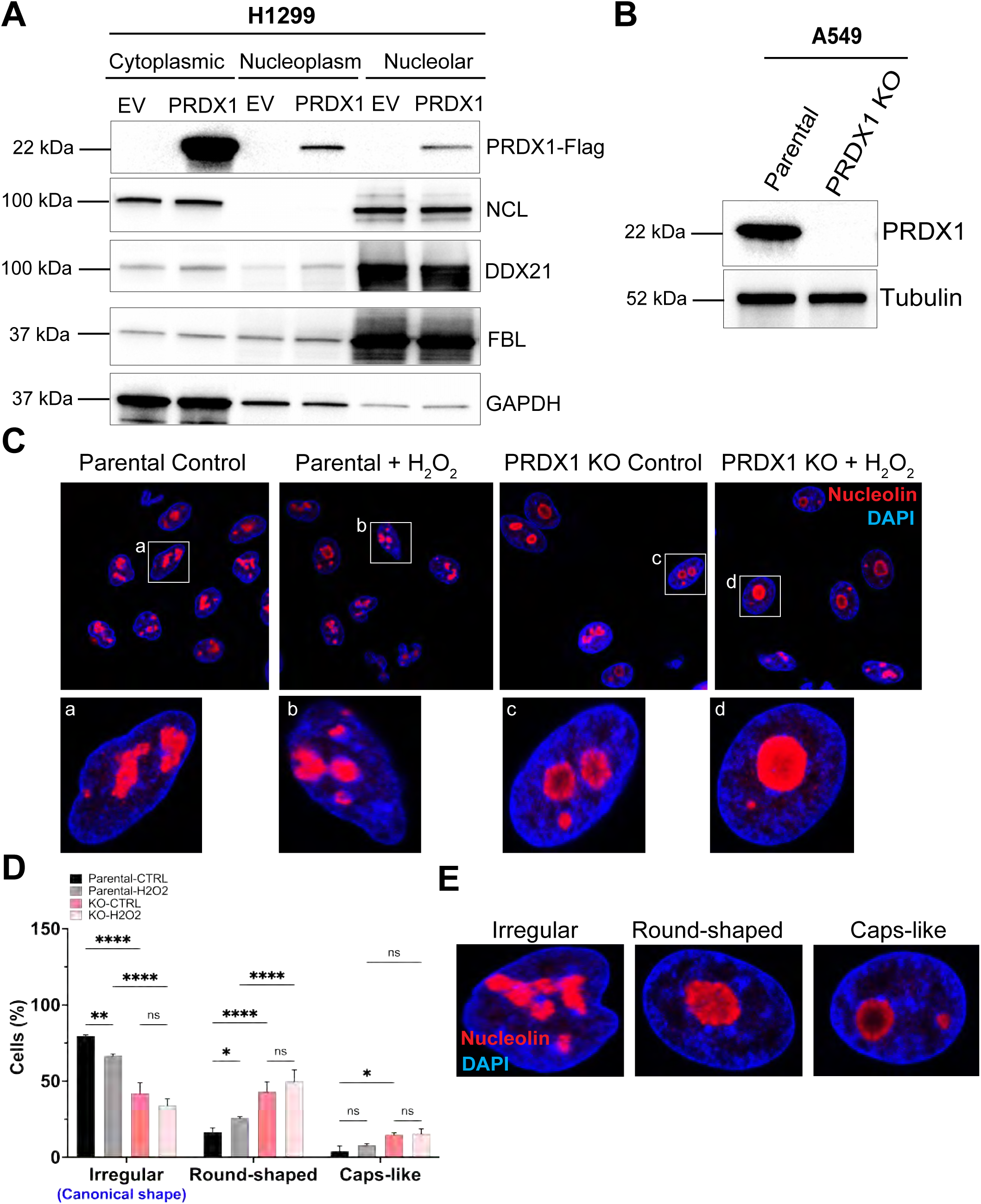
Loss of PRDX1 induces nucleolar abnormalities. (**A**) Western blot analysis of human non–small cell lung cancer H1299 cells transfected with either an empty vector (EV) or a wild-type PRDX1-Flag expression vector (PRDX1-Flag). Cells were transfected 24 hours prior to lysate collection for cellular fractionation. NCL, DDX21, and FBL serve as nucleolar protein markers, while GAPDH indicates enrichment of cytosolic and nucleoplasmic fractions. (**B**) Western blot analysis of human non–small cell lung cancer A549 parental cells and PRDX1 KO cells. Tubulin was used as loading control. (**C**) Confocal microscopy images of A549 parental cells and PRDX1 knockout (KO) cells. Oxidative stress was induced by treatment with 200 µM H_2_O_2_ for 30 minutes. Nuclei and nucleoli were visualized using DAPI (blue) and an anti-Nucleolin (NCL) antibody (red) respectively. Lower magnification images of each condition and a selected nucleus at higher magnification are shown (a-d). (**D**) Comparison of nucleolar shapes. In each condition, nucleoli were classified as canonical nucleoli (irregular shape), round-shaped, or cap-like shaped, and percentages were compared. Statistical analysis was assessed by two-way ANOVA. *P < 0.05, **P < 0.01, **P < 0.0001, ns, not significant. (**E**) Representative images showing the three distinct nucleolar morphologies observed in parental and PRDX1 KO cells.

### PRDX1 loss induces nucleolar genomic instability

Previous reports linked nucleolar abnormalities to nucleolar stress and genomic instability ^1,25,26^. The ribosomal DNA (rDNA) exhibits unique vulnerabilities: its highly repetitive DNA sequence renders it inherently prone to genomic instability; the rDNA loci are subject to replication-transcription conflicts; and the promoter and intergenic spacers regions are enriched in G-rich sequences capable of forming G-quadruplex structures. Furthermore, the elevated metabolic activity within the nucleolus exacerbates these stresses, predisposing rDNA to double-strand breaks (DSBs) ^1,25,26^. We therefore analyzed nucleolar abnormalities in PRDX1 knockout cells to determine if they play a role in nucleolar genomic stability. To elucidate whether the enzymatic activity of PRDX1 is required for nucleolar homeostasis, we utilized parental A549 cells (PAR), PRDX1 knockout cells (KO), and PRDX1 KO cells complemented with either wild-type PRDX1 (Res) or a mutated form of PRDX1 that is devoid of peroxidatic activity (Mut). The mutant form is characterized by mutations in two essential thiol residues (the peroxidative Cp-SH and the resolving Cr-SH) of PRDX1 (Fig. S1A). Using cell fractionation, we confirmed the nucleolar localization of the three genetic variants by cofractionation with nucleolar markers, including NCL, Nucleophosmin (NPM1) and DDX21, as well as a poor enrichment in GAPDH (Fig. S1B). We then stained the cells to detect R-loops using the S9.6 antibody. The level of nucleolar R-loops increased by 50% in PRDX1 KO cells and was significantly reversed in PRDX1 Res cells (Fig 2A, and B). In contrast, PRDX1 Mut cells exhibited R-loop levels close to that observed in PRDX1 KO cells (Fig 2A, and B). As R-loop formation often coincides with the stabilization of G-quadruplexes (G4s), we examined G4s using the BG4 antibody. A similar pattern of nuclear G4 signal was observed in PRDX1-deficient cells, with high levels in the KO and Mut (Fig. 2C and D). As R-loop and G4 stabilization have been implicated in the generation of transcription-replication conflicts, we examined the localization of the Fanconi Anemia complex member FANCD2 that localizes to stalled forks and sites of R-loop-mediated damage ^27^. We observed a clear elevation in FANCD2 in PRDX1 KO cells, an effect that was again rescued in PRDX1-Res, but not Mut cells (Fig. 2E and F). To determine if the stabilization of DNA secondary structures was accompanied by nucleolar DNA breaks, we measured γ-H2AX levels and observed an overall marked increase in γ-H2AX foci in both PRDX1 KO and Mut cells, while parental and PRDX1-Res cells exhibited very low levels of γ-H2AX foci (Fig. 2G and H). Also, the nucleolar γ-H2AX levels in PRDX1 KO and Mut cells were significantly higher than that of parental and PRDX1-Res cells. Altogether, these findings indicated that the enzymatic activity of PRDX1 was essential for both nucleoplasmic and nucleolar genomic stability, implying a key role for PRDX1 in preventing endogenous DNA damage and maintaining nucleolar homeostasis.

**Figure 2:**
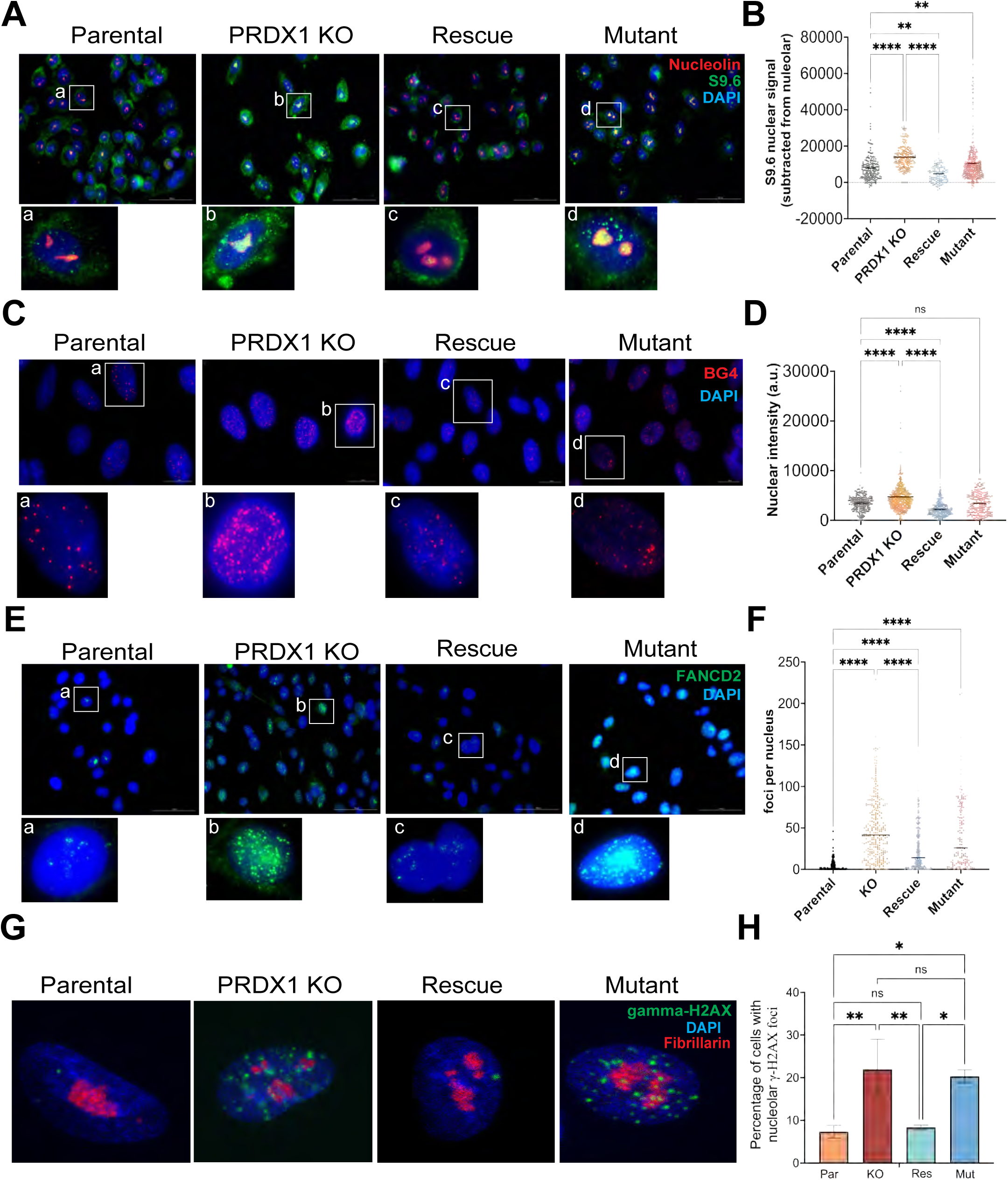
Loss of PRDX1 induces both nucleoplasmic and nucleolar genomic instability. (**A**) Images of A549 cells costained with S9.6 for anti–R-loops detection (green), anti-Nucleolin (red), and DAPI (blue) and analyzed by immunofluorescence as described in *Methods*. A representative image of independent experiments is shown. At least 100 cells were scored for each condition. (**B**) Quantification of images shown in (A). Statistical significance was assessed by two-way ANOVA. **P < 0.01, ***P < 0.001. (**C**) Cells costained with anti-G4 (red), and DAPI (blue) and revealed by immunofluorescence. A representative image of independent experiments is shown. At least 100 cells were scored for each condition. (**D**) Quantification of images shown in (C). Statistical significance was assessed by two-way ANOVA. ****P < 0.0001, ns, not significant. (**E**) Cells were costained with anti-FANCD2 (green), and DAPI (blue), and revealed by immunofluorescence. A representative image of independent experiments is shown. At least 100 cells were scored for each condition. (**F**) Quantification of images shown in (E). Statistical significance was assessed by two-way ANOVA. ****P < 0.0001. (**G**) Cells costained with anti-gamma-H2AX (green), anti-Fibrillarin (red), and DAPI (blue) and analyzed using immunofluorescence. A representative image of independent experiments is shown. At least 100 cells were scored for each condition. (**H**) Quantification of images shown in (G). Statistical significance was assessed by two-way ANOVA. *P < 0.05, **P < 0.01, ns, not significant. Note. For each set of staining (A-F), lower magnification images of each condition and a selected nucleus at higher magnification are shown (a-d).

### PRDX1 is essential for rDNA transcription and ribosome biogenesis

Since PRDX1 loss induced the activation of key hallmarks of nucleolar genomic instability, we sought to determine whether PRDX1 loss may alter transcription activity in the nucleolus. To directly assess the impact of PRDX1 KO on RNA polymerase-I (POL-I) occupancy at the rDNA unit, including the transcribed regions and the intergenic regions (IGS) (Fig. 3A), we employed chromatin immunoprecipitation (ChIP) followed by quantitivative real time PCR (qRT-PCR) in parental A549 and PRDX1 KO cells. The analysis revealed that PRDX1 KO resulted in reduced POL-I binding to rDNA coding regions and lower active transcription in the same regions, as revealed by POL-I and H3K4me3 enrichment at the rDNA loci respectively (Fig. 3B, and C). Treatement with H_2_O_2_ to induce oxidative stress further disminished POL-I occupancy at the transcribed rDNA loci, including the regions encoding the 18S and 28S ribosomal RNA. The specificity of POL-I occupancy was confirmed by the lack of its binding to the IGS. It is worth noting that the IGS regions are poised for active transcription as previously reported ^28^, consistent with the strong enrichment of H3K4me3 observed across this region (Fig 3.C). The partial reduction in POL-I binding to rDNA coding regions in PRDX1 KO cells was mitigated in PRDX1 Res cells, while the effect was not mitigated in PRDX1 Mut cells (Fig. S2A). These data indicated that PRDX1 loss impairs POL-I transcriptional activity, an effect which is further enhanced by exposure to hydrogen peroxide.

**Figure 3:**
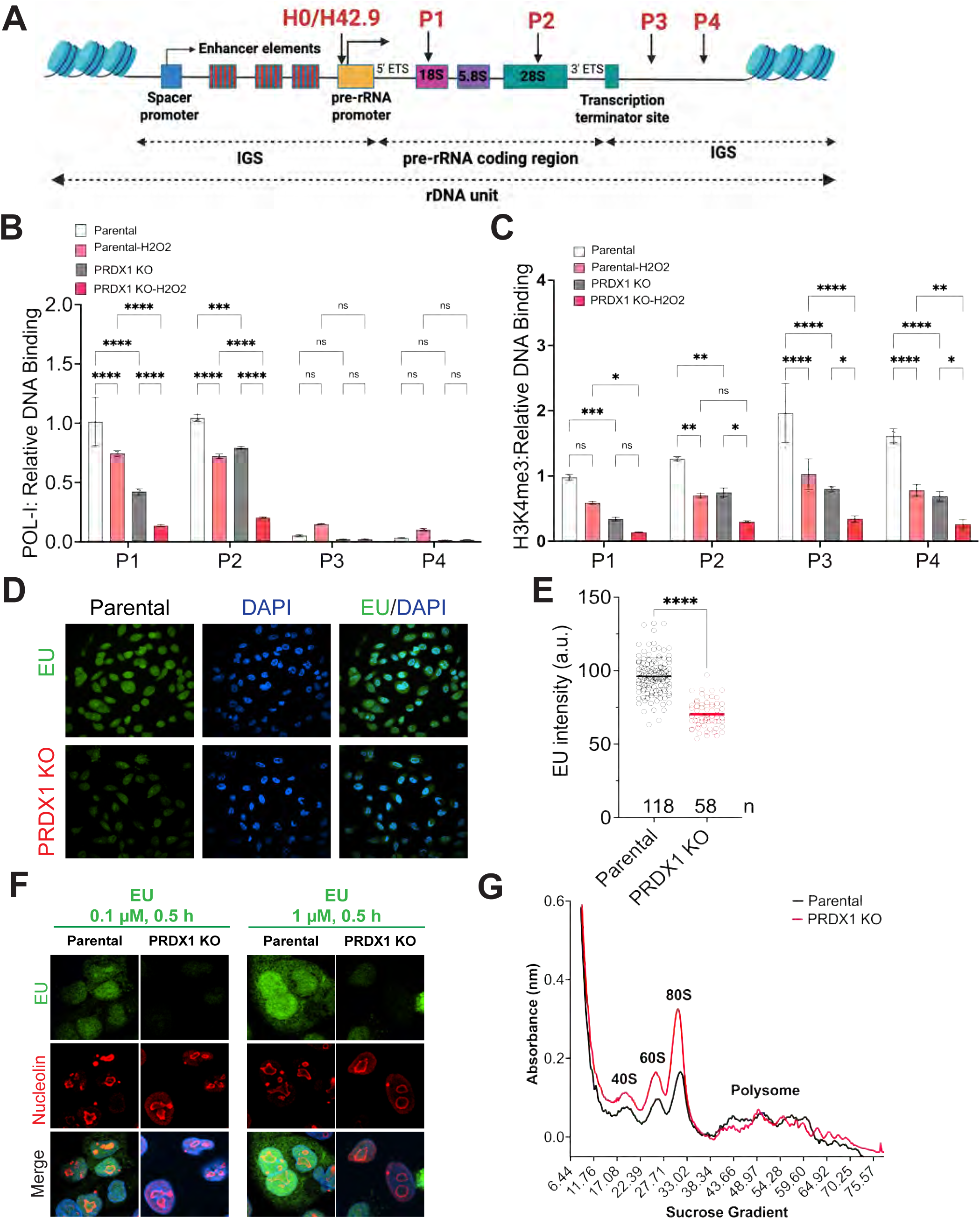
Loss of PRDX1 alters chromatin structure of rDNA and ribosome biogenesis. (**A**) Schematic diagram of the human ribosomal DNA (rDNA) unit and primers used for qPCR. Primer sets include H0 and H42.9 for the promoter region, P1 for 18S ribosomal RNA (rRNA), P2 for 28S rRNA, and P3/P4 for the intergenic spacer (IGS) region. Primers sequence can be found in Materials and Methods (**B and C**) Chromatin immunoprecipitation (ChIP) assay showing the enrichment of rDNA in RNA Polymerase I (POL-I) (**B**), and Histone H3 trimethyl-Lysine 4 (H3K4me3) (**C**) immunoprecipitates. Statistical analysis was performed using two-way ANOVA. *P < 0.05, **P < 0.01, ***P < 0.001, ****P < 0.0001, ns, not significant. (**D-F**) Representative fluorescence microscopy images showing EU (green) incorporation in A549 parental and *PRDX1* KO cells. DAPI was used to counterstain nuclei (**D**). (**E**) Statistical analysis of EU intensity was performed using two-way ANOVA (****P < 0.0001). (**F**) Representative images showing EU (green) and Nucleolin (red) staining in A549 parental and *PRDX1* KO cells. Nucleolin serves as a nucleolar marker, and DAPI (blue) was used for nuclear counterstaining. The overlap between EU and Nucleolin signals indicates sites of rRNA synthesis. (G) Polysome profiling in A549 parental and PRDX1 KO cells.

To determine whether reduced POL-I occupancy translates into functional defects in ribosome biogenesis, we analyzed rRNA synthesis and processing in PRDX1 knockout cells (Fig. S3A). First, we assessed nascent RNA synthesis in single-cells by pulse-label click chemistry with 5-ethyl uridine (EU). Consistent with reduced POL-I binding (Fig. 3B), EU incorporation was strongly reduced in PRDX1 deficient cells (Fig. 3D and E) suggesting a global reduction in RNA synthesis. This reduction also reflects the fact that rRNA constitutes the major RNA species within the cell. Remarkably, loss of PRDX1 was accompanied by depletion of the total levels of nascent rRNA, as revealed by colocalization of EU and NCL (Fig. 3F). Second, we examined the extent to which PRDX1 loss impacted rRNA processing. Using Northern blot analysis with [32P]-orthophosphate labeled ITS1 probe to study steady-state level of rRNA species, we observed only modest changes in global rRNA precursors (Fig. S3B, and C). However, downstream intermediate precursors, such as 21S and 18S-E, were diminished in PRDX1 knockout cells (Fig. S3B, and C). This slight reduction may reflect a delay in pre-RNA transcription and further processing of late stages of small ribosomal subunit maturation when PRDX1 is depleted. Because rRNA synthesis is intrinsincally coupled to ribosome biogenesis, we analyzed the extent to which PRDX1 loss affects ribosomal assembly and translation using a sucrose gradient assay for ribosome profiling. PRDX1 loss led to a drastic increase in the monosome fractions (40S, 60S, and 80S mRNA bound with single ribosome), while slightly decreasing polysome fractions (mRNA bound with multiple ribosomes) (Figure 3G). Altogether, these data support a role for PRDX1 in facilitating rDNA transcription, rRNA synthesis and processing, subsequent ribosome assembly, and ultimately translational capacity.

### PRDX1 loss induces a nucleolar DNA damage response and recruitment of MRN complex at rDNA loci

A previous report demonstrated that nucleolar DNA damage led to MRN recruitment to the nucleolus, with MRN specifically binding the rDNA regions, and repression of POL-I mediated transcription ^7^. ATM-mediated phosphorylation of TCOF1, a nucleolar factor implicated in ribosome biogenesis, promoted MRN recruitment to damaged rDNA sites, even when lesions occurred at distal locations from the nucleolus, a phenomon known as trans-inhibition of POL-I transcription ^7^. Although the underlying mechanism of ATM activation remains unclear, we hypothesized that oxidative stress arising from PRDX1 loss may trigger ATM activation via its auto-phosporylation and subsequent recruitment of MRN complex to the rDNA through interactions with TCOF1. To test this, we performed cell fractionation in PRDX1 KO A549 cells and assessed the protein levels of phospho-ATM, MRN and TCOF1. PRDX1 KO cells displayed heightened ATM phosphorylation across fractions, particularly in the nucleoplasmic and nucleolar fractions, consistent with the presence of chronic oxidative stress in these cells (Fig. 4A). The nucleoplasmic activation of ATM was likely reflective of widespread DNA damage in PRDX1-deficient cells, as previously demonstrated ^16,19–21^. ATM activation was accompanied by a slight nucleolar accumulation of the MRN subunit NBS1, coupled with a massive elevation of nucleolar TCOF1 in PRDX1 KO cells (Fig. 4A). To rule out cell line specific effects, we depleted PRDX1 in H1299 cells using a short hairpin RNA interference (shRNA) approach. Similar to the phenotypes observed in A549 cells, PRDX1 knockdown led to ATM activation and elevated levels of TCOF1 in the nucleolus (Fig. 4B). Immunofluorescence imaging confirmed a marked increase in TCOF1 foci within the nucleolus of PRDX1 KO cells, an effect further exacerbated by treatment with hydrogen peroxide (Fig. 4C, and D), while parental and unstressed cells did not exhibit strong and distinct TCOF1 foci, consistent with previous reports ^10^.

**Figure 4:**
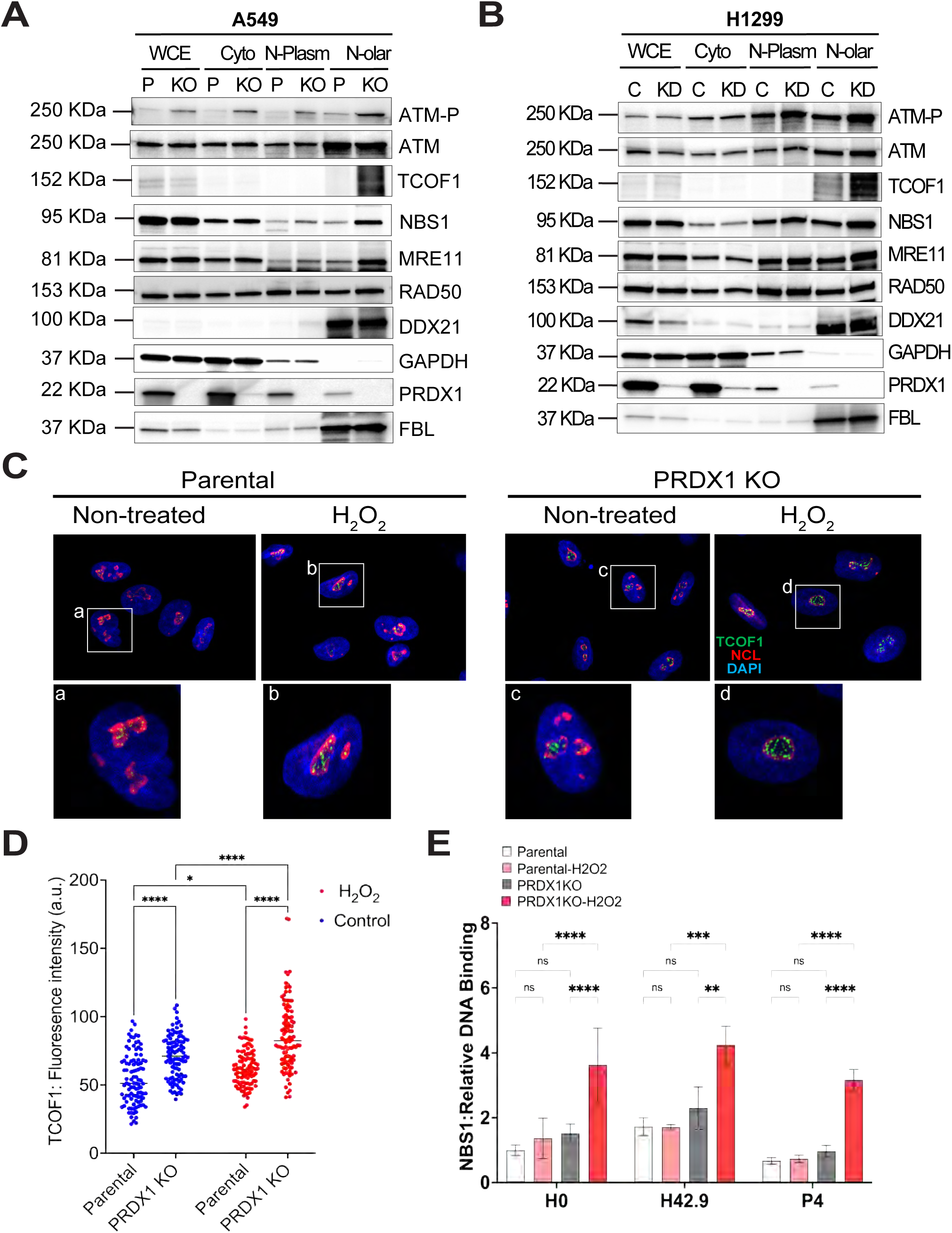
Loss of PRDX1 induces nucleolar DNA damage response via MRN recruitment and NBS1 enrichment at the rDNA loci. (**A, B**) Western blot analysis of human non–small cell lung cancer A549 parental cells and PRDX1 KO cells (**A**) and human non–small cell lung cancer H1299 control and cells infected with short hairpin RNA interference targeting PRDX1 (**B**), following cell fractionation. Fibrillarin (FBL), and DDX21 serve as nucleolar protein markers, while GAPDH indicates enrichment of cytosolic and nucleoplasmic fractions. Note: P, parental, KO, PRDX1 KO, C, control shRNA, KD, shPRDX1. WCE, whole cell extract; Cyto, cytoplasmic fraction; N-Plasm, nucleoplasmic fraction; N-olar, nucleolar fraction. (**C**) Representative confocal microscopy images of A549 parental and PRDX1 KO cells stained with antibodies against TCOF1 (green) and Nucleolin (NCL) (red) with or without H_2_O_2_ treatment (200µM, 30min). Lower magnification images of each condition and a selected nucleus at higher magnification are shown (a-d). (**D**) Quantification of TCOF1 signal intensity per nucleus. (**E**) ChIP assay of A549 parental and PRDX1 KO cells using an anti-NBS1 antibody. The primer sets H0 and H42.9 are located in the promoter region, and P4 is in the intergenic spacer region (see Figure 3A). Statistical analysis was performed using two-way ANOVA. *P < 0.05, **P < 0.01, ***P < 0.001, ****P < 0.0001, ns; not significant.

We next examined whether ATM activation in PRDX1-deficient cells was coupled with the recruitment of the MRN subunit NBS1 to rDNA upon oxidative stress. Parental and PRDX1-KO A549 cells were treated with H₂O₂ for 30 min, followed by ChIP to assess NBS1 occupancy at rDNA loci. For the quantitative real-time PCR, we utilized two sets of primers mapping a promoter region 5’of the transcription start site (H0 and H42.9), and one primer pair mapping a segment inside the intergenic spacer (P4), respectively, as previously employed (Fig. 3A) ^1,7^. PRDX1 KO cells exhibited a threefold increase in NBS1 occupancy both at the rDNA promoter regions and the intergenic spacer after H₂O₂ exposure (Fig. 4E), implying a widespread recruitment of MRN complex across the rDNA units. PRDX1 KO marginally altered POL-I occupancy at rDNA promoter regions (Fig. S4A), an effect that was not significantly amplified by exposure to H₂O₂. These findings align with previous reports showing that DNA damage promotes NBS1 recruitment to rDNA promoters without significantly affecting POL-I binding to the promoter region ^7^. Our results suggest that PRDX1 is required for the nucleolar DNA damage response to oxidative stress and that PRDX1 loss drives the recruitment of MRN to rDNA regions.

To further assess whether the degree of MRN binding to rDNA loci is influenced by the presence of PRDX1 and its redox sensing ability, we examined ROS levels in parental, PRDX1-KO, PRDX1-Res, and Mut cells, upon exposure to H₂O₂, followed by quantification of NBS1 occupancy at rDNA loci using ChIP. PRDX1 KO and Mut cells drastically accumulated ROS, compared to parental and PRDX1 Res cells that had similar basal ROS levels (Fig. 5A). Consistently, PRDX1 KO cells displayed elevated pS1981-ATM and TCOF1 levels in the nucleolus, particularly following exposure to H₂O₂ (Fig. 5B), an effect that was mitigated by ectopic expression of PRDX1 (Fig. 5B). Using ChIP, we observed enhanced NBS1 binding at rDNA loci in PRDX1 KO upon oxidative stress (Fig. 5C), while NBS1 occupancy in PRDX1 Res cells mirrored the levels of parental cells (Fig. 5C). This further implied that PRDX1 is essential for suppressing ROS and preventing aberrant recruitment of NBS1 to rDNA loci. Remarkably, the enhanced NBS1 enrichment at the promoter region in PRDX1 KO cells was strongly associated with repression of POL-I binding at the rDNA coding regions and reversed in PRDX1-Res cells (Fig. 5D). Taken together, these data further underscore the requirement for PRDX1 for mitigating oxidative stress-induced rDNA transcriptional repression potentially preserving ribosome biogenesis and subsequent translational regulation.

**Figure 5:**
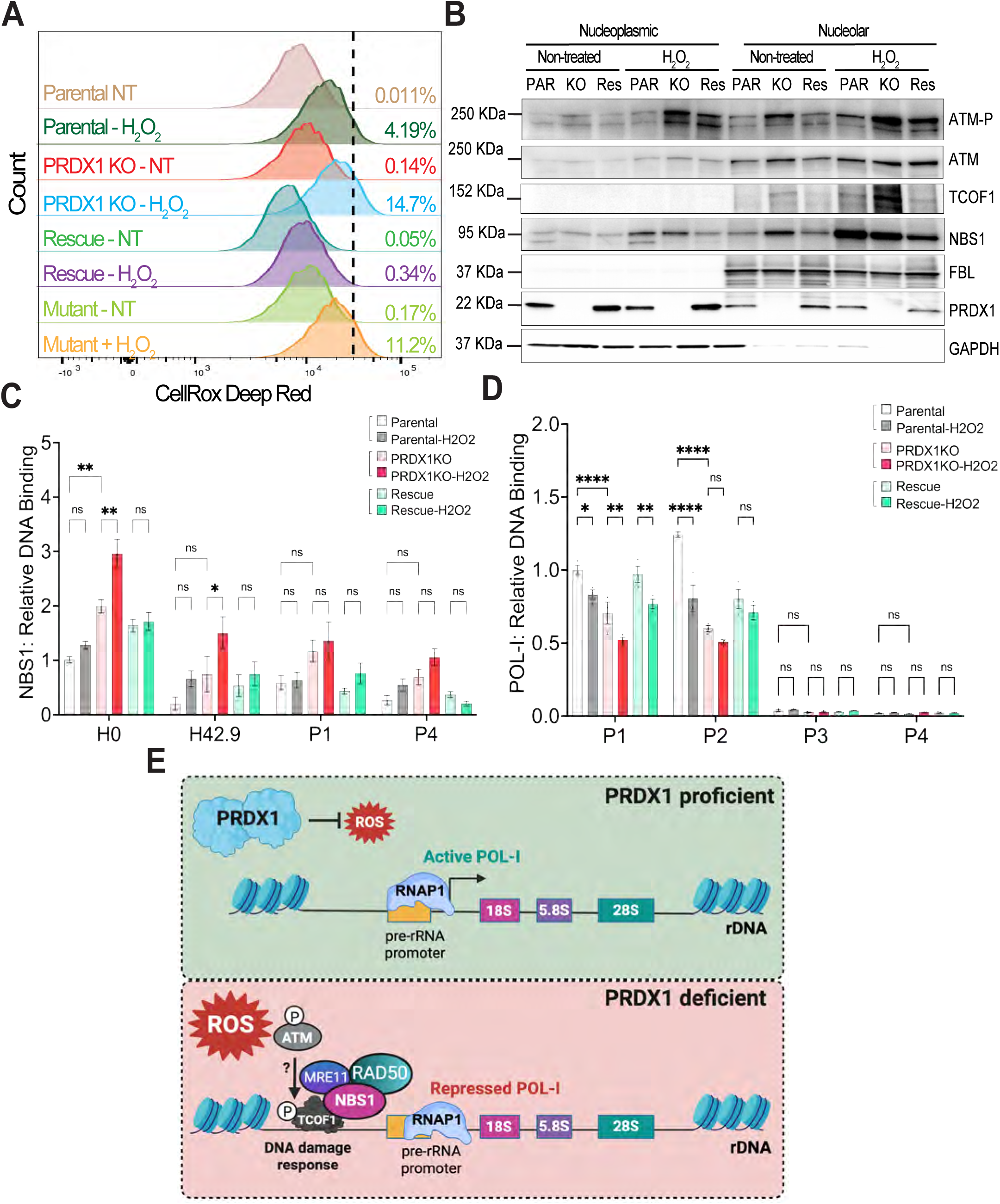
PRDX1 ectopic expression reverses the nucleolar DDR and restores POL-I activity. (**A**) Comparison of ROS levels in A549 parental, PRDX1-KO, PRDX1-KO Rescue (PRDX1-KO complemented with wild type PRDX1), and Mutant (PRDX1-KO complemented with CPRS mutated PRDX1) cells (see Figure S1A) following H_2_O_2_ treatment (200 µM for 30 min). Oxidative stress was detected by *CellRox Deep Red* and monitored by flow cytometry. The response to H_2_O_2_ treatment was compared to the baseline where endogenous reactive oxygen species (ROS) were detected in non-treated (NT) parental cells. Loss of PRDX1 induces more ROS when treated with H_2_O_2_ (PRDX1 KO-H_2_O_2_), while Rescue-H_2_O_2_ cells resist the induction of ROS levels as parental cells. Mutant cells phenocopied PRDX1 KO cells. (**B**) Western blot analysis of human non–small cell lung cancer A549 parental, PRDX1 KO, and PRDX1 rescue cells treated with H_2_O_2_ (200 µM) for 30 min, followed by cell fractionation. Fribrillarin (FBL) serves as nucleolar protein marker, while GAPDH indicates enrichment of cytosolic and nucleoplasmic fractions. Note: Par, parental cells, KO, PRDX1 KO cells, and Res, PRDX1 wild type-rescued cells. (**C**) ChIP assay showing the enrichment of rDNA in NBS1 immunoprecipitates with or without H2O2 treatment (200 µM, 30 min). The primer sets H0 and H42.9 are located in the promoter region, P1 is in the coding region, and P4 is in the intergenic spacer region (see Figure 3A). Note that NBS1 binding was reduced in Rescue cells (PRDX1-KO complemented with wild type PRDX1) similar to parental cells. (**D**) ChIP assay showing the enrichment of rDNA in RNA Polymerase I (POL-I) immunoprecipitates with or without H_2_O_2_ treatment (200 µM, 30 min). Primer sets P1 and P2 are located in 18S and 28S ribosomal RNA, respectively, and P3/P4 are in the intergenic spacer region (see Figure 3A). Note that POL-I binding in Rescue cells returned to levels observed in parental cells. Statistical analysis was performed using two-way ANOVA. *P < 0.05, **P < 0.01, ***P < 0.001, ****P < 0.0001, ns; not significant. **(E)** In PRDX1-proficient cells (green), reactive oxygen species (ROS) are efficiently scavenged to prevent aberrant Pol I-dependent rDNA transcription, thereby maintaining nucleolar homeostasis. Loss of PRDX1 (pink) resulted in accrued ROS level potentially triggering activation of the DNA damage response kinase ATM through autophosphorylation and activation of nucleolar DNA damage response via recruitment and enrichment of the MRN complex, and TCOF1 at rDNA loci, particularly at the pre-rRNA promoter region. Accumulation of NBS1 restricts POL-I-dependent transcription at rDNA coding regions, including 18S, 5.8S, and 28S, potentially delaying rRNA synthesis and protecting the nucleolar genome from further oxidative damage. Elevated DNA breaks caused by ROS accumulation following PRDX1 loss may trigger a comparable response.

In summary, we show that PRDX1 functions as a key redox factor that maintains nucleolar homeostasis and genomic stability. Under oxidative stress, PRDX1 knockout cells rapidly accumulate MRN complex proteins and TCOF1 in the nucleolus, followed by NBS1 enrichment at rDNA loci. This nucleolar DNA damage response is strongly associated with repression of POL-I-dependent transcriptional activity (Fig. 5E). It remains unclear whether this nucleolar DDR is mediated by ATM activation.

## Discussion

In this study, we identified PRDX1 as an essential regulator of nucleolar homeostasis, safeguarding rDNA from oxidative stress and inappropriate POL-I-dependent transcription of damaged templates. Cells depleted of PRDX1 exhibited aberrant nucleolar morphology indicative of nucleolar stress. Earlier reports from our group and others demonstrated that PRDX1 loss is sufficient to induce DNA damage, oxidative nuclear lesions and replicative stress ^16,19–21^. Here, we show that PRDX1 functions as a key redox factor maintaining nucleolar homeostasis and genomic stability. Under oxidative stress, PRDX1 knockout cells rapidily accumulate MRN complex in rDNA loci, as reflected by NBS1 enrichment in ChIP assay, likely limiting rDNA transcription. Because PRDX1 loss creates a state of chronic oxidative stress, NBS1 recruitment and TCOF1 activation are likely mediated through ATM activation as previously reported ^7^. Indeed, NBS1 recruitment at the rDNA promoter coincides with reduced POL-1 occupancy at rDNA coding regions following H_2_O_2_ treatment, particularly in PRDX1 knockout cells, suggesting a DDR-driven surveillance mechanism. These results are consistent with studies showing that ATM signalling suppresses local POL-I transcription in response to chromatin breaks ^8^. We propose that PRDX1 loss induces ATM-driven DNA damage responses in both the nucleoplasmic and nucleolar compartments. This response is further amplified by exposure to oxidative stress ^7^. More specifically, the presence of NBS1 and TCOF1 in the nucleolus likely endows PRDX1 knockout cells with the ability to rapidily respond to nucleolar stress through enhanced NBS1 recruitment to rDNA loci and attenuation of POL-I activity across the coding regions upon oxidative stress. Other reports have highlighted the importance of NBS1 in the oxidative stress-induced DNA damage response, further strengthening our findings ^7,10,29–31^. NBS1 nucleolar activity has also been linked to DNA end processing for the repair of oxidative stress-induced DNA lesions ^9^. Changes in NBS1 binding in response to oxidative stress are consistent with our ROS data showing that the lack of PRDX1 favors ROS accumulation, potentially rendering cells more vulnerable to elevated stress and DNA damage-induced transcription repression. However, it remains unclear whether elevated ROS levels in PRDX1 knockout cells directly trigger ATM phosphorylation to initiate MRN complex recruitment, or whether ROS-induced nucleolar DNA breaks serve as the primary signal activating the DDR. Nonetheless, these findings highlight the essential role of PRDX1 in mitigating nucleolar stress and preventing alterations in transcriptional activity.

In PRDX1-deficient cells, we observed elevated nucleolar DNA breaks, as well as the accumulation of DNA structures, including R-loops and G4s, that have been associated with transcription-replication conflicts and elevated genome instability. These defects were reversed by expression of ectopic PRDX1, but not in cells expressing PRDX1 Mut. The stabilization of these DNA structures likely reflects chronic oxidative stress and are consistent with the perturbation of POL-I-dependent transcription in PRDX1-deficient cells. These observations are reminiscent of models of chronic oxidative stress, including ATM loss-associated secondary DNA structures, as previously reported ^32^. While our findings link PRDX1 to nucleolar genomic stability, it remains unclear whether PRDX2, which has well-established nuclear functions ^33,34^, also contributes to nucleolar homeostasis and ribosome biogenesis, warranting further investigation. Likewise, our current findings do not rule out the possibility of widespread DNA damage in PRDX1-deficient cells, including in the nucleoplasm, that could contribute to trans-repression of POL-I-dependent transcription. This implies that nucleolar DNA damage alone may not fully account for the observed perturbations in rDNA transcriptional homeostasis.

Although reduced POL-I transcription suggests a global effect on rRNA synthesis, as confirmed by EU incorporation assay, we observed only modest changes in rRNA processing. Northern blot analysis revealed unchanged steady-state levels of early rRNA precursors (45S, 41S, 30S and 26S) but significantly reduced late-stage intermediate precursors such as 21S and 18S-E, consistent with oxidative stress-mediated defects in rRNA processing ^35^. These steady-state RNA levels may reflect a balance between synthesis and processing rates. The unchanged abundance of early precursors, despite reduced transcription, indicates a “processing bottleneck” that compensates for decreased synthesis. Impairment of late-stage processing slows the conversion of early intermediates into mature rRNA species, therby maintaining their steady-state levels under dimished transcriptional input. The pronounced reduction in 21S and 18S-E intermediates demonstrates that PRDX1 loss causes a dual defect: reduced POL-I dependent transcription coupled with impaired late stage maturation of the small ribosomal subunit. This processing defect ultimately results in downstream translational dysfunction revealed by the polysome profiling. Moreover, the observed increase in monosome fractions coupled with reduced polysome fractions links PRDX1 deficiency to abnormal ribosomal assembly, consistent with other models of nucleolar genomic instability and perturbed ribosome biogenesis ^1,36^.

Our findings highlight two major advances. First, we establish that PRDX1, canonically characterized as a redox signaling enzyme, exerts a critical nuclear function in maintaining nucleolar genomic stability. Second, because the nucleolar genome is the powerhouse of ribosome biogenesis and subsequent protein synthesis, PRDX1’s role extends beyond ROS scavenging to encompass the regulation of translational capacity and potentially impacting cell fate. Future studies should determine the extent to which PRDX1 influences global protein homeostasis and the integrated stress response. Collectively, our work identifies PRDX1 as a major regulator of nucleolar homeostasis, providing new insights into how the nucleolar genome is protected from oxidative stress-induced damage.

## MATERIALS AND METHODS

### Cell culture

Cells were grown at 37°C and 5% CO_2_ in a humidified incubator. A549-Cas9 cells were grown and maintained in Dulbecco’s Modified Eagle Medium (DMEM) (Fisher Scientific) supplemented with 10% Fetal Bovine Serum (Thermo Scientific) and 2 μg/mL blasticidin for maintaining Cas9 expression. NCI-H1299 cells were grown in RPMI 1640 medium (Fisher Scientific) supplemented with 10% Fetal Bovine Serum (Thermo Scientific) and 1 mM sodium pyruvate (Gibco). All media used in this study were supplemented with penicillin and streptomycin (Gibco). A549-Cas9 PRDX1 knockout cells were generated using CRISPR/Cas9 genome editing. LentiGuide-Puro (Addgene, #52963) was digested with BsmBI-v2 (NEB). sgPRDX1-1(5’-GCGCTTCGGGTCTGATACCAA-3’), and sgPRDX1 -3(5’- TGAAAGCAATGATCTCCGTG-3’) guide RNA oligos targeting human PRDX1, were cloned into the lentiviral plasmid and further utilized for the production of lentiviral particles where sgPRDX1-1 and sgPRDX1-3 plasmids were co-transfected with lentiviral packaging plasmids psPAX2 and pCMV-VSVG into HEK-293 FT cells using Lipofectamine® 3000 (Thermo Scientific). At 48 hours post-transfection, the culture medium was collected and filtered through a 0.45 μM filter to be incubated with A549-Cas9 cells in the presence of polybrene (8 μg/ml). At 72 hours post-infection, infected cells were selected with 1 μg/mL puromycin for 7 days. Single-cell clones were expanded and screened by Western Blot for protein levels of PRDX1. Cells with ectopic expression of wild-type PRDX1 and mutation of both C_p_-SH and C_R_-SH (Cys52/Cys172) of PRDX1 were used as mentioned in ^21^. NCI-H1299 PRDX1 knockdown cells were generated by infection similar to the method mentioned above, with lentiviral particles produced using shRNA expressing plasmids (pLKO.1) targeting human PRDX1 (TRCN0000029513). Cells were selected with 0.5 μg/mL puromycin for 7 days and then maintained in the medium with 0.25 μg/mL puromycin.

### Transfection

For plasmid transfection in NCI-H1299, 7 million cells were seeded in a T75 culture flask the day before transfection. On the following day, 4 μg of plasmid (pcDNA3.1-Flag or pcDNA3.1-PRDX1-flag) was transfected into the cells with jetOPTIMUS® DNA transfection Reagent (VWR, Cat. 76299-634) following the manufacturer’s protocol.

### Flow cytometry

For reactive oxygen species measurement, cells were incubated with the HBSS buffer (Gibco) with 5 μM CellRox Red (Invitrogen) for 30 min at 37°C after indicated drug treatment. After incubation, cells were PBS-washed three times and trypsinized. The data was acquired using a BD LSRFortessa SORPII and analyzed witg FlowJo.

### Immunofluorescence and image quantification

Cells were seeded at a density of 50,000 cells per chamber of an 8 well chambered slide. Cells were washed with phosphate-buffered saline (PBS), fixed in 4% paraformaldehyde (Sigma-Aldrich) for 30 min at room temperature, then blocked in PBS with 0.05% Tween 20 +0.1% Triton X-1000 for 1 hour at room temperature and stained with primary antibody, Cell Signaling Technology-NCL (E5M7K) and FBL (C13C3) (1:400) overnight at 4°C. The next day cells were washed with phosphate-buffered saline (PBS) followed by secondary antibodies, Thermo Fisher Scientific-Alexa Fluor 488 and Alexa Fluor 594 (1:400) in the same buffer at room temperature for 2 hrs. Quantification was performed on at least 100 cells.

For assessing co localization, a confocal Leica Stellaris STED microscope at ×63 magnification 1.4 NA (with magnification to a resolution of approximately 40 nm per pixel) was used.

### R-loop staining using S9.6 antibody

Cells were seeded at a density of 50,000 cells per coverslip in a 24 well plate and 24hrs later fixed using ice cold Methanol for 20 minutes and quenched using 1M Glycine at room temperature for 10 min. Cells were permeabilized using ice cold acetone for 10 min. Following this cells were treated with 1:200 dilutions of RNase T1 (Thermo Fisher Scientific, EN0541), ShortCut RNase III (New England Biolabs, M0245S) at 37°C for 1hr across all conditions to minimize dsRNA or RNA secondary structures. For negative control cells were treated with 20U/ml of RNaseH1 (New England Biolabs, M0297S) at 37°C for 2hrs. Cells were blocked using 1% BSA with 0.1% triton solution made in 1X PBS for 1hr at room temperature. Following this, cells were incubated with primary antibodies S9.6 for R-loops (Millipore MABE1095) at 1:250 dilution and nucelolin (ab136649) overnight at 4°C. Finally cells were incubated with respective secondary antibodies for 1hr at room temperaturend. Cells were stained with DAPI and mounted using Vectashield antifade mounting media. Imaging was done at 60X magnification using the Ziess Microscope. For data analysis Gen5 software was used. The nucelolar signal marked by anti-nucelolin antibody was used to subtract the nucleolar R-loop signal from the total nuclear intensity.

### G-quadruplex staining using BG4 antibody

Cells were seeded at a density of 50,000 cells per coverslip in a 24 well plate. 24hrs later cells were fixed using 4% PFA. Cells were permeablilized using 0.1% triton for 10 minutes followed by RNase A treatment (0.25ug/ml) for 1hr at 37°C. Cells were then fixed with blocking solution for 1hr at 37°C (0.5% Goat serum with 0.1% triton in 1X PBS). Primary antibody, BG4 (homemade) was added in 1:5000 dilution in blocking buffer for 1hr at 37°C. Following primary antibody, cells were incubated with anti-FLAG antibody (CST D6W5B) at 1:1000 dilution in blocking buffer for 1hr at 37°C. Finally, cells were incubated with anti-rabbit secondary antibody at 1:1000 dilution in blocking buffer for 1hr at 37°C. Cells were stained with DAPI and mounted for imaging.

### EU labeling and quantification of signal intensity in immunofluorescence assays

Cells were seeded at a density of 50,000 cells per chamber in an 8-well chamber (Nunc™ Lab-Tek™ II CC2™, Thermo Fisher, #154941), and labeled with 0.5 mM EU for 1 hour. Click-iT RNA imaging Kit (Invitrogen C10329) was used for EU detection. The quantification of signal intensity for EU and antibodies was done using Fiji (ImageJ) software. A uniform circular region defined by DAPI staining was consistently applied to all isogenic cell line samples for quantification of mean signal intensities. The resulting values were used to construct individual signal profiles. Processed data were subsequently imported into GraphPad Prism 10 (GraphPad Software Inc.) for visualization.

### Cell Fractionation

After respective treatments, 12 M cells were collected and resuspended in 600 μL hypotonic buffer (10 mM HEPES [pH 7.9], 10 mM KCl, 1.5 mM MgCl2, 0.5 mM DTT). Cells were homogenized using syringe with needle for 8 times. After centrifugation (500 g) for 10 min, the supernatant was recovered to serve as the cytoplasmic fraction. The pellet was resuspended in 300 μL S1 buffer (0.25 M sucrose, 10 mM MgCl2), overlaid onto 300 μL S2 buffer, and centrifuged (1,500 g) for 5 min. The pellet was resuspended in 300 μL S2 buffer (0.35 M sucrose, 0.5 mM MgCl2) to serve as the nuclear fraction. The nuclear fraction was then sonicated (30 s sonication, 60 s intervals for 5 cycles, 50% amplitude) and overlaid onto 300 μL S3 buffer (0.88 M sucrose, 0.05 mM MgCl2), and centrifuged (3,000 g) for 10 min. The supernatant was collected as the nucleoplasmic fraction. The pellet was resuspended in RIPA buffer to serve as the nucleolar fraction. Phosphatase inhibitor cocktail 3 (Sigma-Aldrich) and cOmplete ULTRA EDTA-Free Protease inhibitors (Roche) were added to all fraction samples. The fractions were resolved on SDS-PAGE gels and blotted.

### Protein extraction and Western blotting

Cells were lysed on ice in RIPA Lysis buffer supplemented with phosphatase inhibitor cocktail 3 (Sigma-Aldrich) and cOmplete ULTRA EDTA-Free Protease inhibitors (Roche). Cells were sonicated for 15s at 50% (Fisherbrand) and spun down for 5 min at 14,000 rpm at 4°C. Supernatants were quantified using the Bradford assay (Thermofisher Scientific). Protein lysates (30 μg) were resuspended in Laemmli’s buffer supplemented with β-mercaptoethanol (Sigma-Aldrich, 1:10). Proteins from cellular extracts were separated by electrophoretic migration using Tris-Glycine-SDS buffer (Bio-Rad). Semidry transfer was performed with a TransBlot Turbo apparatus (Bio-Rad) in TransBlot Turbo transfer buffer (Bio-Rad) for 30 min at 25 V. TransBlot Turbo mid-size nitrocellulose membranes were blocked for at least 1 hour in 0.1% Tween 20 and 5% BSA and incubated with primary antibodies overnight at 4°C. Visualization was performed using West Femto ECL (Thermo Fisher Scientific) substrate. Images were acquired using a ChemiDoc imaging system (Bio-Rad).

The following antibodies were used at 1:1000 dilution from Cell Signaling Technology-PRDX1 (D5G12), p95/NBS1(D6J5I), NCL (E5M7K), Phospho-ATM (Ser1981) (D25E5), ATM (D2E2), Mre11 (31H4), Rad50 (E3I8K), and at 1:2000 dilution for FBL (C13C3), DDX21, NPM1 (E7W4P). 1:2500 for GAPDH-HRP (14C10), α-Tubulin (11H10), Vinculin (E1E9V). Anti-TCOF1 was purchased from Sigma-Aldrich.

### Preparation of Northern blotting probes and Northern blot

Synthetic DNA oligonucleotide, Human ITS1 (GGCCTCGCCCTCCGGGCTCCGTTAATGAT) was ordered from Integrated DNA Technologies (IDT) and resuspended in dH_2_O. 1 μl of the 6 μM DNA was added to 2 μl of [^32^P]-γ-ATP (3000 Ci/mmol) (Revvity), 1 μl of 10× T4 PNK buffer, 1 μl of T4 PNK (NEB) and 14 μl of dH_2_O for 1 h at 37°C. The reaction was brought up to 100 μl of dH_2_O and the reaction mix was filtered using gel filtration G-25 columns (GE LifeSciences) to remove the unincorporated nucleotides.

RNA was extracted with PureLink™ RNA Mini Kit (Invitrogen) according to manufacturer’s instructions. 10μg of extracted RNA was loaded on a denaturing gel, electrophoresed and passively transferred on a Hybond N^+^ membrane (Amersham) in 20× SSC buffer [Saline-Sodium Citrate Buffer, Cepham Lifesciences]. Membrane was pre-hybridized in 10 ml of ULTRAhyb-Oligo hybridization buffer (Invitrogen) for 1 h at 60°C. Pre-hybridization solution was removed, and fresh ULTRAhyb-Oligo hybridization buffer was added to the membrane containing 15μl of the end-labeled ITS1 probe. The membrane was then incubated for 1 h at 60°C followed by overnight hybridization at 37°C. The following day the membrane was washed with 2× SSC/ 0.1% SDS at 40°C and exposed to film. Typhoon FLA9500 scanner (GE Healthcare) was used to acquire the radioactive signal.

### Chromatin immunoprecipitation

All ChIP experiments were performed using the Pierce Magnetic ChIP kit (Thermo Fisher Scientific, 26157) according to the manufacturer’s instructions. Briefly, 1.2 x 10^7^ cells were seeded on a 150 mm dish. For H_2_O_2_ treatment, cells were incubated with 200 µM H_2_O_2_ for 30 minutes. Cells were cross-linked with a 1% formaldehyde solution for 10 minutes at room temperature, followed by treatment with 0.125 M glycine for 5 minutes. Fixed cells were lysed, and the chromatin was digested using 2 units of micrococcal nuclease (MNase) for 15 minutes at 37°C. Chromatin prepared from 4 x 10^6^ cells was incubated with 10 µg of antibody or normal rabbit IgG as a control. The following antibodies were used for ChIP reactions: Anti-NBS1 (Novus Biologicals, NB100-143), Anti-POLR1A (Cell Signaling Technology, 24799), and Anti-H3K4me3 (Cell Signaling Technology, 9751). The Anti-Polymerase II antibody and primers for GAPDH are included in the ChIP kit. Precipitated DNAs were subjected to quantitative real-time PCR (qPCR) to evaluate the enrichment of protein binding. qPCR was carried out using the QuantStudio 5 system (Applied Biosystems) according to the manufacturer’s protocol. PCR products were examined by melting curve analysis, and fold changes were calculated using the 2–ΔΔCT method. The primers used for qPCR are listed below.

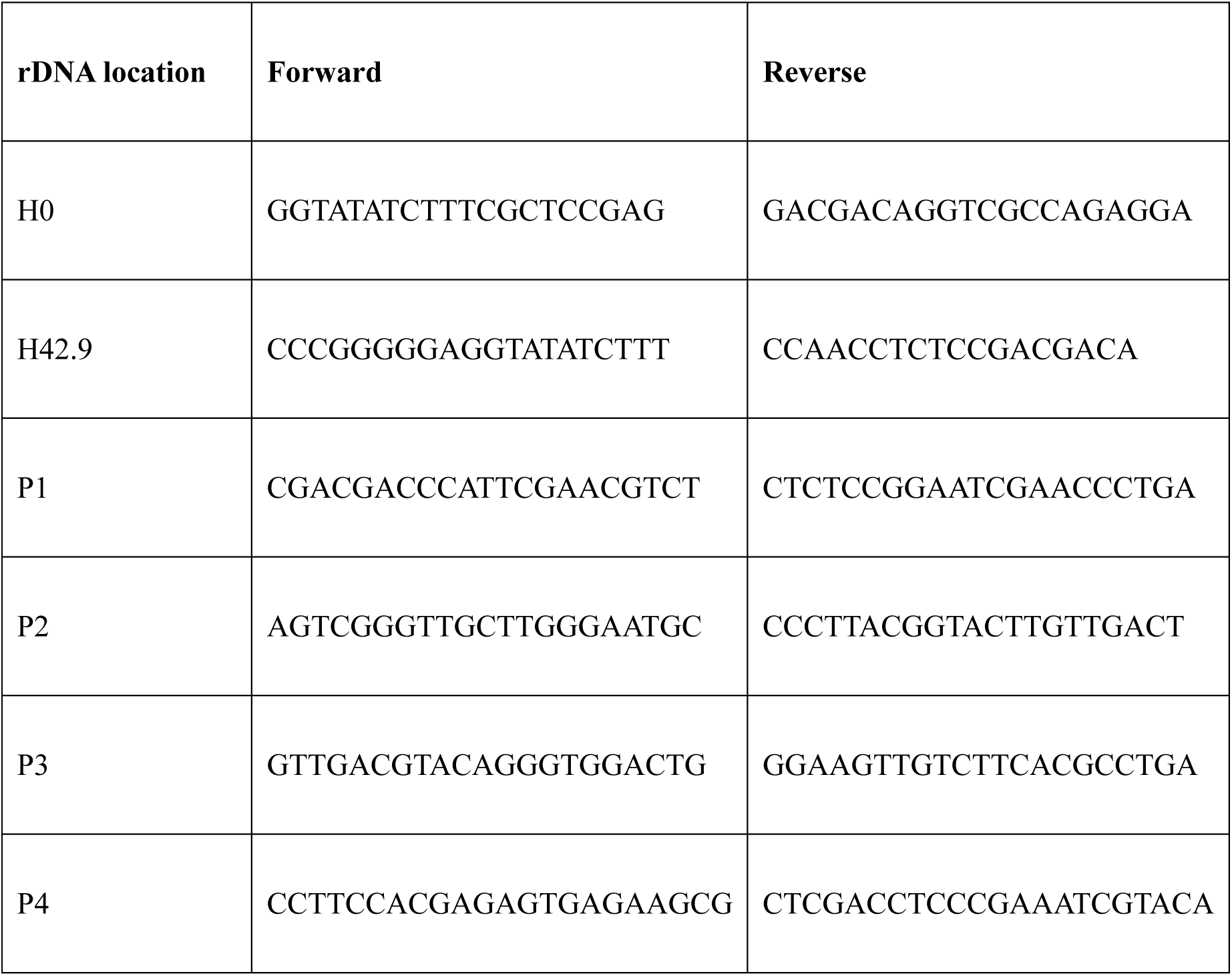

### Polysome profiling

Cells were seeded in 150 mm dishes per condition to have 60-70% confluency on the day of polysome isolation. For H_2_O_2_ treatment, cells were incubated with 200 µM H_2_O_2_ for 30 minutes/2 hours. Cycloheximide (100 μg/ml) was added for 10 min in the medium and cells were washed in ice-cold PBS. Cells were lysed in 50 mM HEPES (pH 7.3), 100 mM KOAc, 15 mM MgOAc, 1 mM dithiothreitol (DTT), cycloheximide (100 μg/ml), and 1% Triton-X. The lysate was clarified by centrifugation at 14000 rpm for 10 min at 4°C, and the RNA was quantified by measuring absorbance A_260._ The clarified lysate was loaded onto a 10-45% sucrose gradient (25 mM HEPES pH 7.3, 100 mM KOAc, 5 mM MgOAc, 1 mM DTT, cycloheximide (100 μg/ml), and supplemented with *cOmplete Protease Inhibitor*) prepared using a BioComp Gradient Master, then centrifuged for 1 hour and 50 minutes at 41,000 rpm at 4°C in a Beckman SW41Ti rotor. The ribosomal fractions were collected and their absorbance at 254 nm (*A*_254_) was measured using the BioComp fractionator to plot the polysome profiles.

### Data reproducibility and statistical analysis

Prism 10 (GraphPad Software Inc.) was used to determine the statistical significance of experiments stated in the figure legends. Specific biological replicate numbers (n) for each experiment can be found in the corresponding figure legends. Statistically significant differences are labeled with one, two, three, or four asterisks if *p* < 0.05, *p* < 0.01, *p* < 0.001, or *p* < 0.0001, respectively.

## Acknowledgments

We thank the NCI-CCR Microscopy Core for help with the confocal microscopy experiments. We also thank Dr. Keli Agama for their help with the Northern blotting assay. This work is supported by the NCI Center for Cancer Research (CCR) and has been funded in whole or in part with Federal funds from the National Cancer Institute, National Institutes of Health. The content of this publication does not necessarily reflect the views or policies of the Department of Health and Human Services, nor does mention of trade names, commercial products, or organizations imply endorsement by the U.S. Government.

## Author contributions

T.F., V.G., and U.W. designed the experiments. T.F., V.G., S.S., F.B., H.L., C.A., and D.T. performed the experiments. T.F., V.G., S.S., H.L., and D.T. analyzed the data. Y.P., S.O., and T.H.S., provided reagents and expertise and substantially reviewed the manuscript. U.W. wrote the manuscript with inputs from all authors.

## RESOURCE AVAILABILITY

Further information and requests for resources, reagents, or data should be directed to and will be fulfilled by the Lead Contact, Urbain Weyemi.

