## Supplementary Materials for "Peroxiredoxin 1 safeguards the nucleolar genome from oxidative damage"

1. Developmental Therapeutics Branch, NCI Center for Cancer Research, National Cancer Institute, National Institutes of Health, 37 Convent Drive, Bethesda, MD 20892, USA. 2. Radiation Oncology Branch, NCI Center for Cancer Research, National Cancer Institute, National Institutes of Health, Bethesda, MD 20892, USA. 3. Laboratory of Receptor Biology and Gene Expression, NCI Center for Cancer Research, National Cancer Institute, National Institutes of Health, Bethesda, MD 20892, USA.

*** Corresponding author:**

Developmental Therapeutics Branch

National Cancer Institute, National Institutes of Health

37 Convent Drive, Bethesda, MD 20892, USA.

### These authors contributed equally to this study

**Figure S1**: **Detection of wild type and mutant PRDX1 across subcellular fractions**

(**A**) Diagram illustrating PRDX1 active residues. Two PRDX1 proteins act cooperatively as a dimer upon oxidation. PRDX1 contains a conserved cysteine residue (Cys 52) in its N-terminal region, known as peroxidatic Cys ($S_P$-H) which senses $H_2O_2$ and undergoes oxidation forming a sulfenic acid. PRDX1 also contains an additional conserved Cys residue in its C-terminus (Cys 173) known as resolving Cys ($S_R$-H). A disulfide structure is formed in response to $H_2O_2$ and consists of the peroxidatic Cys52 residue from one PRDX1 and the resolving Cys172 residue from the other homodimer subunit. (**B**) **Western blot analysis** of human non–small cell lung cancer A549 parental, PRDX1 KO, PRDX1 rescue, and PRDX1 mutant cells, following cell fractionation. Nucleolin (NCL), NPM1, and DDX21 serve as nucleolar protein markers, while GAPDH indicates enrichment of cytosolic and nucleoplasmic fractions. Note: Par, parental cells, KO, PRDX1 KO cells, Res, PRDX1 wild type-rescued cells, and Mut, PRDX1 C52S/C172S mutant cells.

**Figure S2: Re-expression of PRDX1 protein in PRDX1-KO cells facilitates Pol I binding to rDNA.** qPCR analyses of chromatin samples from A549 parental, PRDX1-KO, PRDX1-KO complemented with wild type PRDX1 (Rescue), and PRDX1-KO complemented with CPRS mutated PRDX1 (Mutant). Chromatin samples were immunoprecipitated using either anti-RNA Polymerase I (Pol I) or normal rabbit IgG (non-specific binding control). The relative amount of immunoprecipitated DNA is compared with input DNA and normalized to the P1 value of the parental control. Primer sets P1 and P2 are located in 18S and 28S ribosomal RNA, respectively, and P3/P4 are in the intergenic spacer region (see Figure 3A). Statistical analysis was performed using two-way ANOVA. *$P < 0.05$, **$P < 0.01$, ***$P < 0.001$, ****$P < 0.0001$, ns; not significant.

**Figure S3: Detection of rRNA precursors in parental and PRDX1 KO cells.**

(**A**) Schematic representation of the maturation pathways of human rRNAs. Approximate location of the northern blotting probe ITS1 is noted in red. (**B**) Northern blotting of cellular RNA in parental and PRDX1 KO cells. A total of 10 μg of RNA was resolved on 1 % agarose formaldehyde gel, transferred to Hybond $N^+$ membrane and probed with end-labeled ITS1 northern blotting probe. PRDX1 loss results in reduction in downstream precursors 21S and 18S-E. (**C**) Quantification of rRNA precursors detected in (B). Statistical analysis was performed using two-way ANOVA. *P < 0.05, ***P < 0.001, ns; not significant. Quantification of rRNA precursors was performed by normalizing signal intensity to ethidium bromide (EtBr)-stained 18S and 28S rRNA bands.

**Figure S4: PRDX1 loss slightly reduces RNA Polymerase I binding to the promoter region.** ChIP assay showing the enrichment of rDNA in RNA Polymerase I (POL-I) immunoprecipitates with or without H2O2 treatment (200 μM, 30 min). The primer sets H0 and H42.9 are located in the promoter region, and P4 is in the intergenic spacer region (see Figure 3A). Statistical analysis was performed using two-way ANOVA. *P < 0.05, **P < 0.01, ***P < 0.001, ****P < 0.0001, ns, not significant.

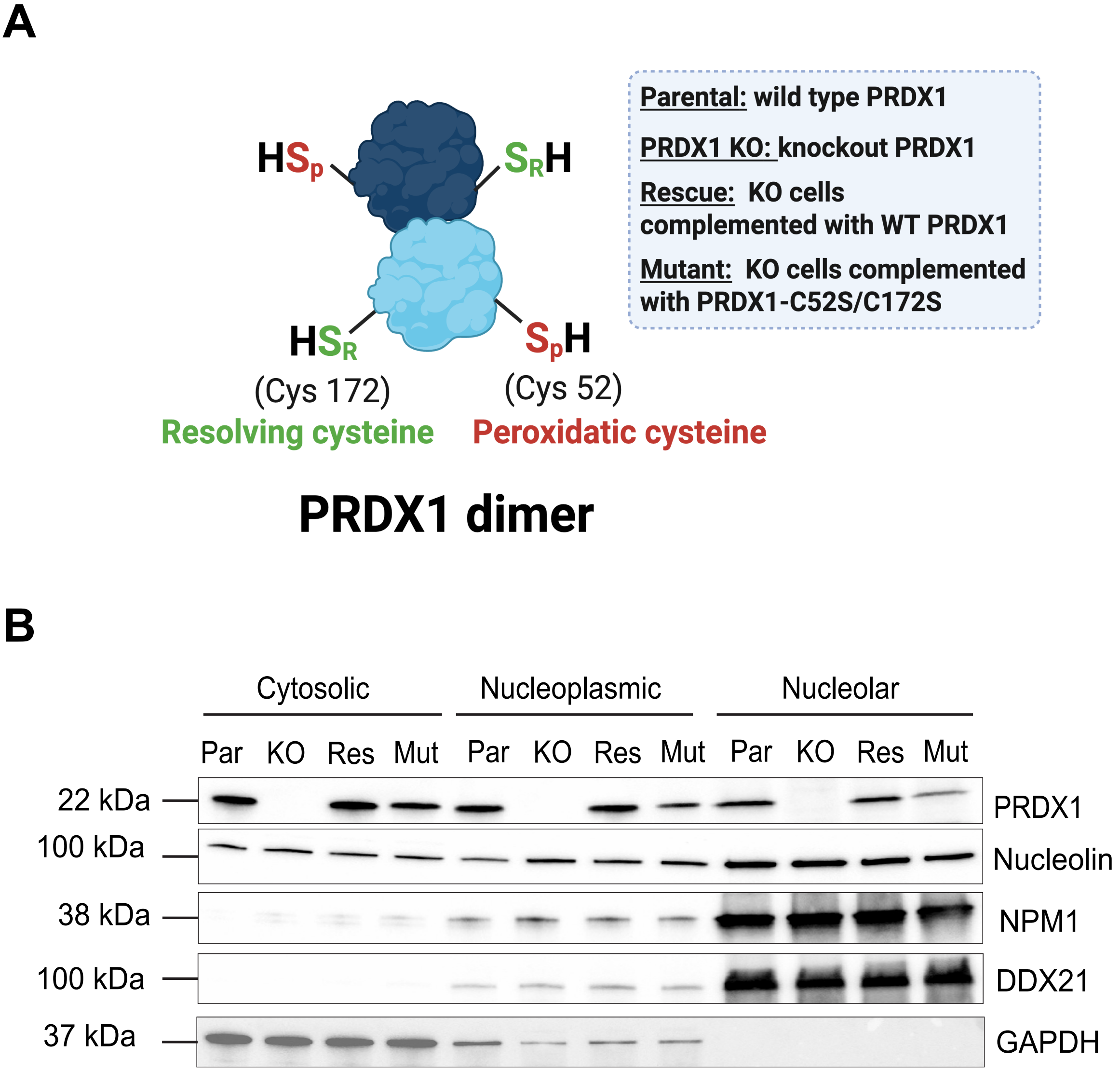
A
HSp
SRH
HSR
SpH
(Cys 172)
Resolving cysteine
(Cys 52)
Peroxidatic cysteine
PRDX1 dimer
Parental: wild type PRDX1
PRDX1 KO: knockout PRDX1
Rescue: KO cells complemented with WT PRDX1
Mutant: KO cells complemented with PRDX1-C52S/C172S
B
Cytosolic
Nucleoplasmic
Nucleolar
Par KO Res Mut Par KO Res Mut Par KO Res Mut
22 kDa
100 kDa
38 kDa
100 kDa
37 kDa
PRDX1
Nucleolin
NPM1
DDX21
GAPDH

Figure S1

A

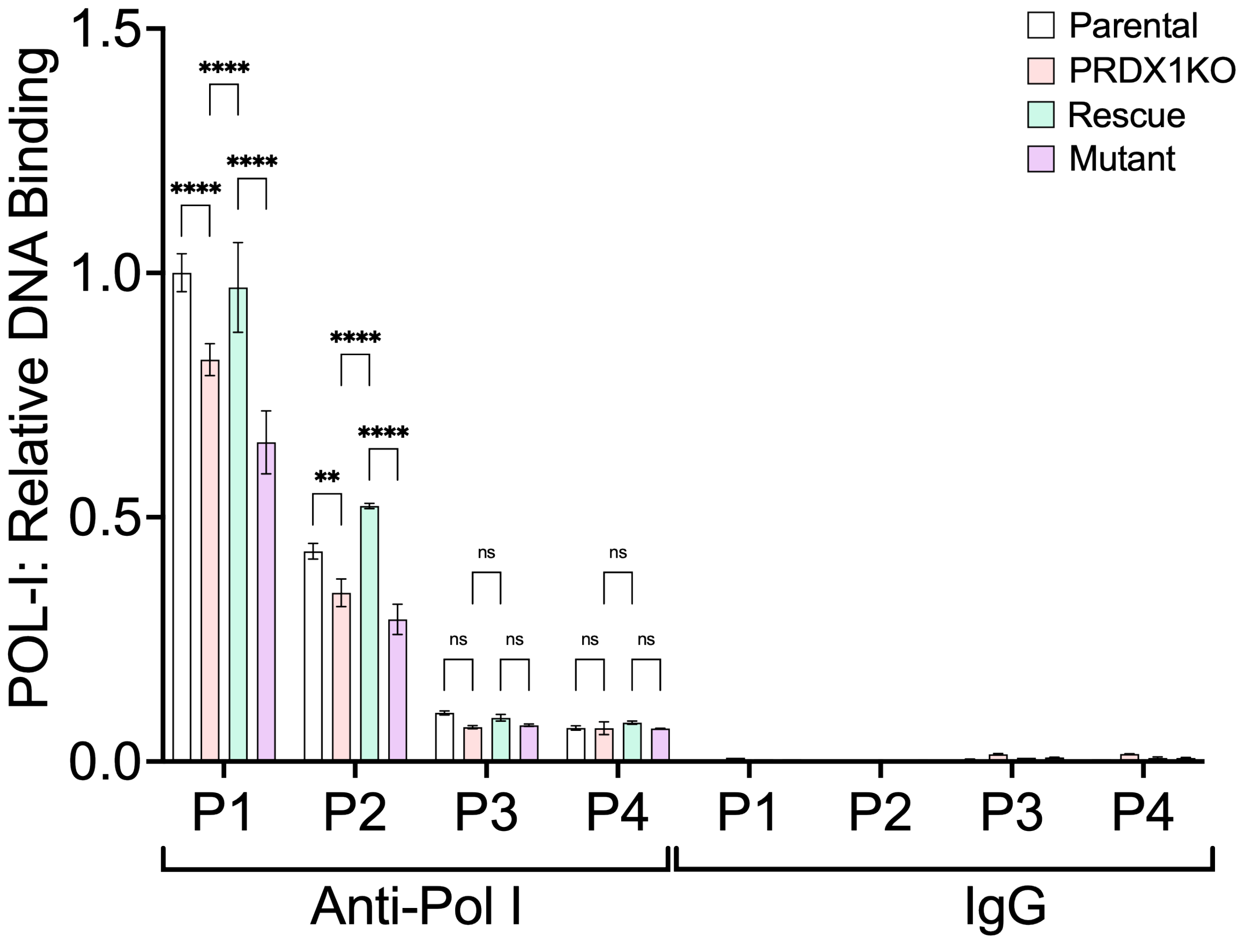

Parental
PRDX1KO
Rescue
Mutant
POL-I: Relative DNA Binding
1.5
1.0
0.5
0.0
****
****
****
****
**
****
ns
ns
ns
ns
ns
ns
P1
P2
P3
P4
P1
P2
P3
P4
Anti-Pol I
IgG


Figure S2

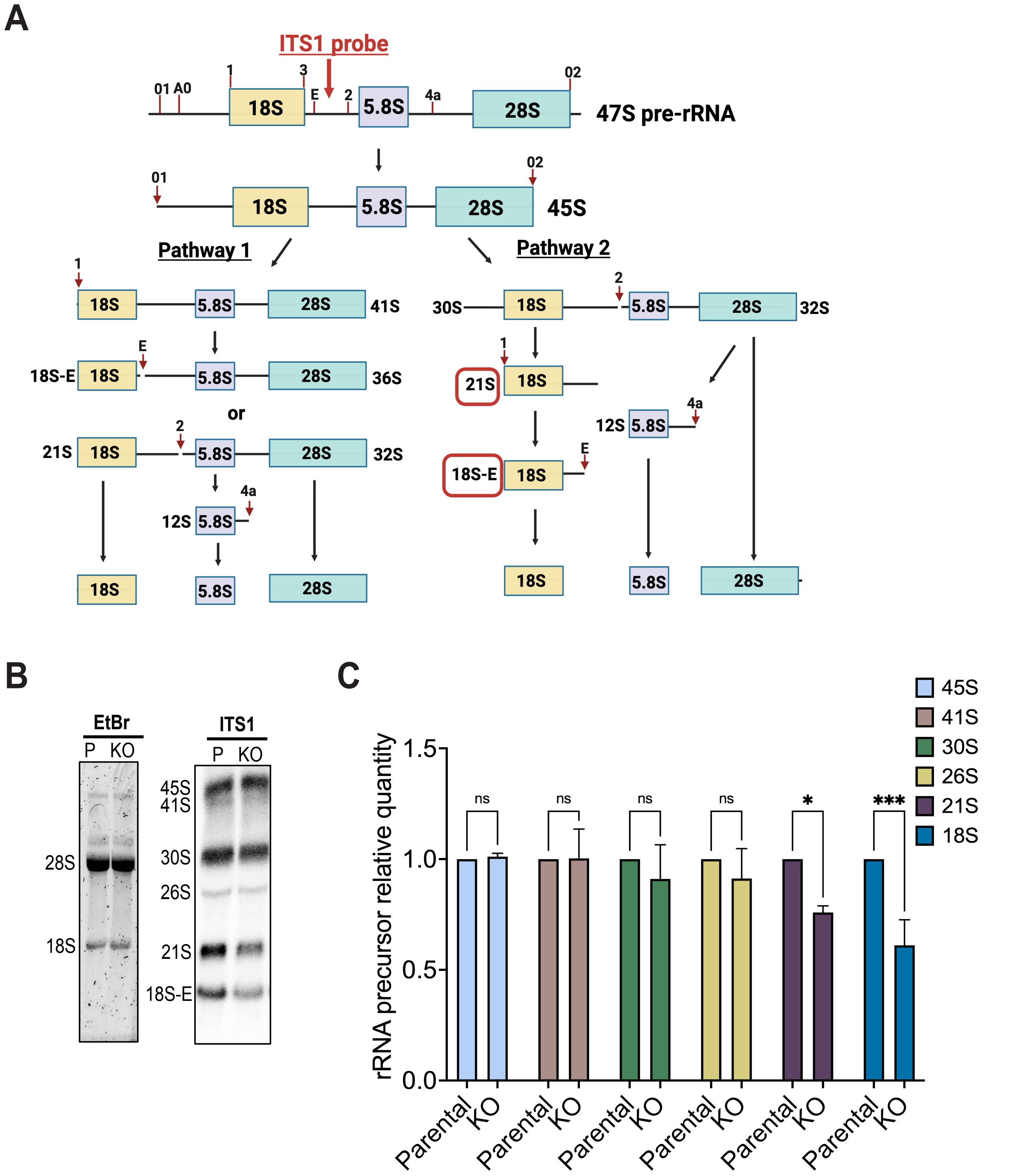
A
ITS1 probe
47S pre-rRNA
45S
Pathway 1
Pathway 2
18S-E
or
21S
12S
41S
36S
32S
30S
B
EtBr
P KO
28S
18S
ITS1
45S
41S
30S
26S
21S
C
rRNA precursor relative quantity
Parental
KO
ns
*
***

Figure S3

A

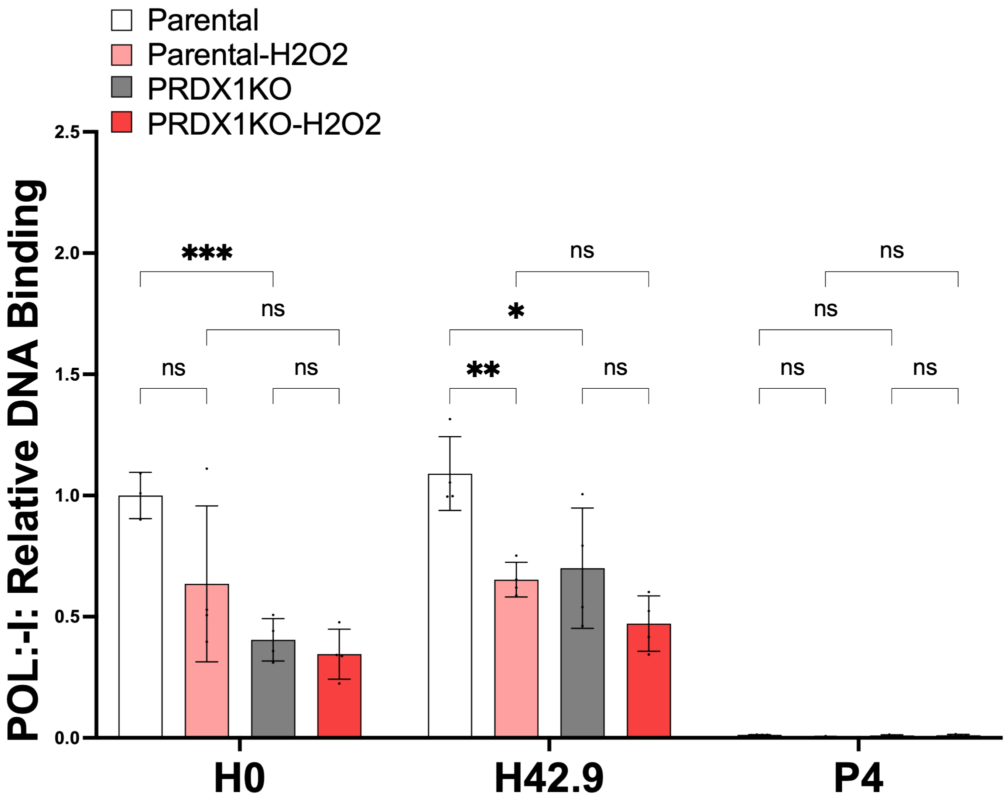
Parental
Parental-H2O2
PRDX1KO
PRDX1KO-H2O2
POL:-I: Relative DNA Binding
2.5
2.0
1.5
1.0
0.5
0.0
***
ns
ns
ns
*
**
ns
ns
ns
ns
ns
ns
H0
H42.9
P4

Figure S4
